# Multimodal Reorganization of Gut–Brain Physiology During the Cephalic Phase of Digestion

**DOI:** 10.64898/2026.11.07.765106

**Authors:** Yasmine M. Kehnemouyi, Todd P. Coleman

**Affiliations:** Department of Bioengineering, Stanford University, Stanford, CA, USA

## Abstract

The vagus nerve plays a central role in gut–brain communication, yet the physiological dynamics associated with cholinergic vagal signaling remain difficult to characterize non-invasively in humans. The cephalic phase response, which prepares the gastrointestinal system for food ingestion through anticipatory vagal pathways, offers a unique opportunity to investigate these interactions. Here, we used simultaneous recordings of cortical, gastric, cardiac, and swallow physiology in healthy adults to characterize brain–body responses during food anticipation and subsequent ingestion. Food-related sensory cues increased gastric myoelectric activity as well as swallow and heart rate while eliciting frequency-specific changes in cortical oscillations. Strikingly, anticipation was accompanied by opposing changes in physiological coordination: synchronization within cortical and gastric networks decreased, whereas coupling between the brain and stomach increased. These changes largely returned toward baseline following ingestion, suggesting a distinct anticipatory organization of brain–body activity. Moreover, gastric responses were associated with various aspects of subjective appetite, whereas cortical responses were selectively associated with craving. Both gastric and cortical responses independently explained the variation in participants’ self-reported craving. Together, these findings reveal that food anticipation reorganizes physiological coordination within and across central and peripheral systems, establishing the cephalic phase as a non-invasive framework for investigating brain–body interactions associated with cholinergic vagal signaling in humans.

## 2 Introduction

The gut–brain axis comprises a complex communication network linking the gastrointestinal system and the brain through neural, endocrine, immune, and metabolic pathways [1]. The vagus nerve is a major neural component of this network, mediating bidirectional communication between the gastrointestinal tract and the central nervous system through afferent sensory pathways and cholinergic efferent signaling [1, 2, 3]. These interactions are essential for normal physiology, with gut-derived signals such as glucagon-like peptide-1 (GLP-1) regulating appetite and metabolism partly through vagal pathways [4, 5]. Disruptions in gut–brain signaling have also been implicated in neurological and psychiatric disorders, including Parkinson’s disease [6, 7]. Despite the importance of these pathways, cholinergic vagal signaling to the gastrointestinal tract remains difficult to characterize non-invasively in humans.

The cephalic phase response offers a unique opportunity to investigate cholinergic vagal signaling in humans by engaging anticipatory digestive pathways before food is consumed. Food-related sensory cues, including the sight, smell, taste, or thought of food, recruit cortical networks involved in sensory processing, attention, and motivation [8, 9]. These cortical responses are accompanied by descending autonomic and motor output, including cholinergic vagal efferent signaling originating from the dorsal motor nucleus of the vagus (DMV), which contributes to preparing the gastrointestinal system for incoming nutrients [8, 10, 3]. This anticipatory preparation manifests as changes in gastric activity, swallow, salivation, and cardiovascular physiology before ingestion begins. Although the cephalic response has a long history in experimental physiology, its potential as a paradigm for understanding coordinated brain–body dynamics remains largely unexplored.

Human studies of cephalic phase responses have traditionally focused on peripheral outputs, including gastric acid secretion, insulin, and pancreatic polypeptide, often using sham-feeding paradigms [8, 11, 12]. In parallel, studies of brain responses to food cues have identified changes in cortical theta, alpha, and beta oscillations associated with sensory, attentional, and motivational processing [13, 14, 15]. Although these approaches have established important features of anticipatory physiology, they have largely characterized central and peripheral responses separately. Whether food anticipation simply recruits these systems in parallel or reorganizes how their activity is coordinated therefore remains unknown. Furthermore, it remains unclear whether cortical and gastric responses reflect shared or distinct aspects of subjective appetite, including hunger, satiety, and craving.

To address these questions, we simultaneously recorded cortical, gastric, cardiac, and swallow physiology in healthy adults during fasting, food anticipation elicited by visual and olfactory cues, and meal ingestion. Food anticipation elicited widespread changes in central and peripheral physiology, accompanied by a striking dissociation between within-system synchronization and cross-system coupling. While synchronization within cortical and gastric networks decreased, coupling between the brain and stomach increased, suggesting that anticipation reorganizes physiological coordination across organizational scales. These changes largely returned toward baseline following ingestion. Moreover, gastric responses were associated with hunger, satiety, and craving, whereas cortical responses were selectively associated with craving, with both independently contributing to individual differences in craving. Together, these findings reveal that food anticipation does not simply activate the brain and digestive system in parallel, but reorganizes their physiological interactions before nutrients enter the gastrointestinal tract. More broadly, this work establishes the cephalic phase as a non-invasive paradigm for investigating physiological responses associated with cholinergic vagal signaling in humans, providing a framework for understanding gut–brain communication and its potential disruption in neurological and gastrointestinal disorders.

## 3 Results

We simultaneously recorded electroencephalography (EEG), electrogastrography (EGG), electrocardiography (ECG), and swallow electromyography (EMG) in 40 healthy adults (mean age, 25.7 *±* 2.5 years) during 15-minute fasted baseline and cephalic phases. Of these, 32 participants also completed the 15-minute ingestion phase following consumption of a personalized study meal; the remaining eight either elected not to consume the meal or were unable to complete the phase because of time constraints. Usable EEG and EGG recordings were available for all 40 participants during baseline and the cephalic phase, whereas data availability during the ingestion phase and across other modalities varied because of phase completion and signal quality. Phase- and modality-specific sample sizes are summarized in Table 1.

**Table 1:**
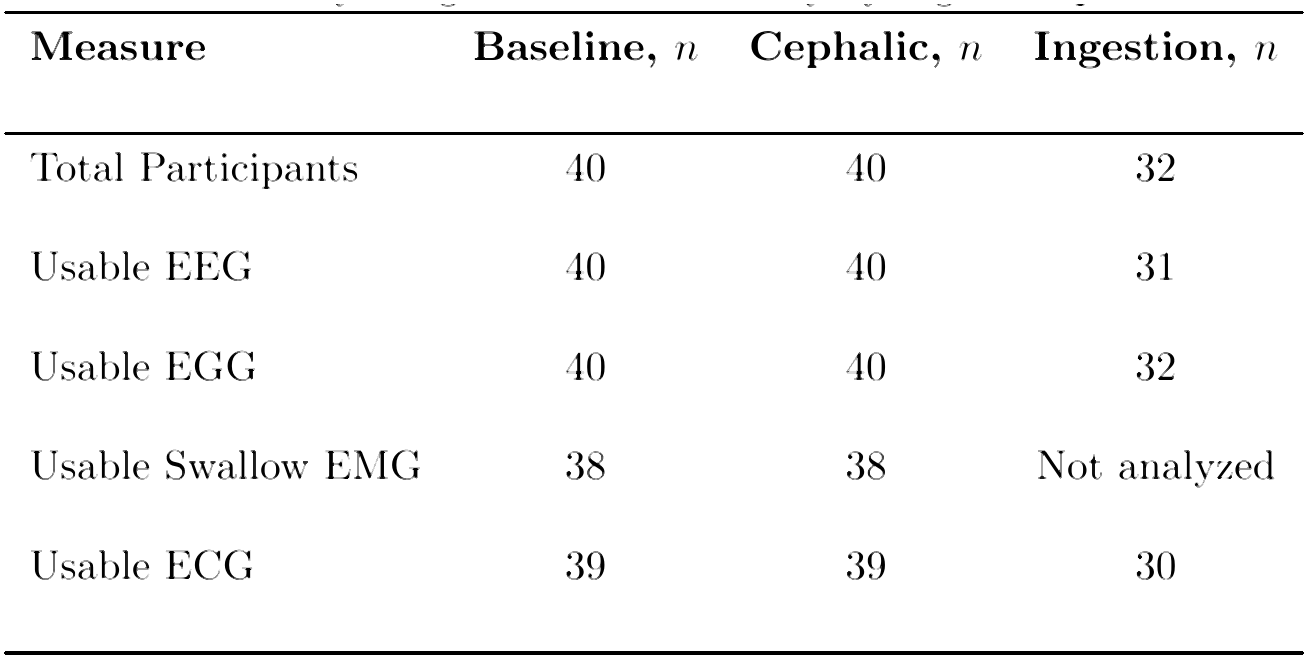
Physiological data availability by digestion phase.

### 3.1 Visceral physiology is modulated during the cephalic phase and evolves following ingestion

To characterize peripheral physiological responses to food anticipation, we quantified swallow activity, heart rate, and gastric myoelectric power within the 0.03–0.07 Hz frequency range (Fig. 1). During the cephalic phase, heart rate increased progressively following food cue presentation, with a significantly greater mean heart rate than baseline (*t*(38) = 5.95, *p <* 0.001, Cohen’s *d_z_* = 0.95; Fig. 1A,B). Swallow activity increased rapidly and remained elevated throughout food anticipation, with a significantly greater mean swallow rate than baseline (paired *t*-test, *t*(37) = 8.63, *p <* 0.001, Cohen’s *d_z_* = 1.40; Fig. 1C,D). Similarly, gastric-band power increased following food cue presentation and remained elevated throughout the cephalic phase, with significantly greater mean power than baseline (*n* = 40, *t*(39) = 6.94, *p <* 0.001, Cohen’s *d_z_* = 1.10; Fig. 1E,F).

**Figure 1:**
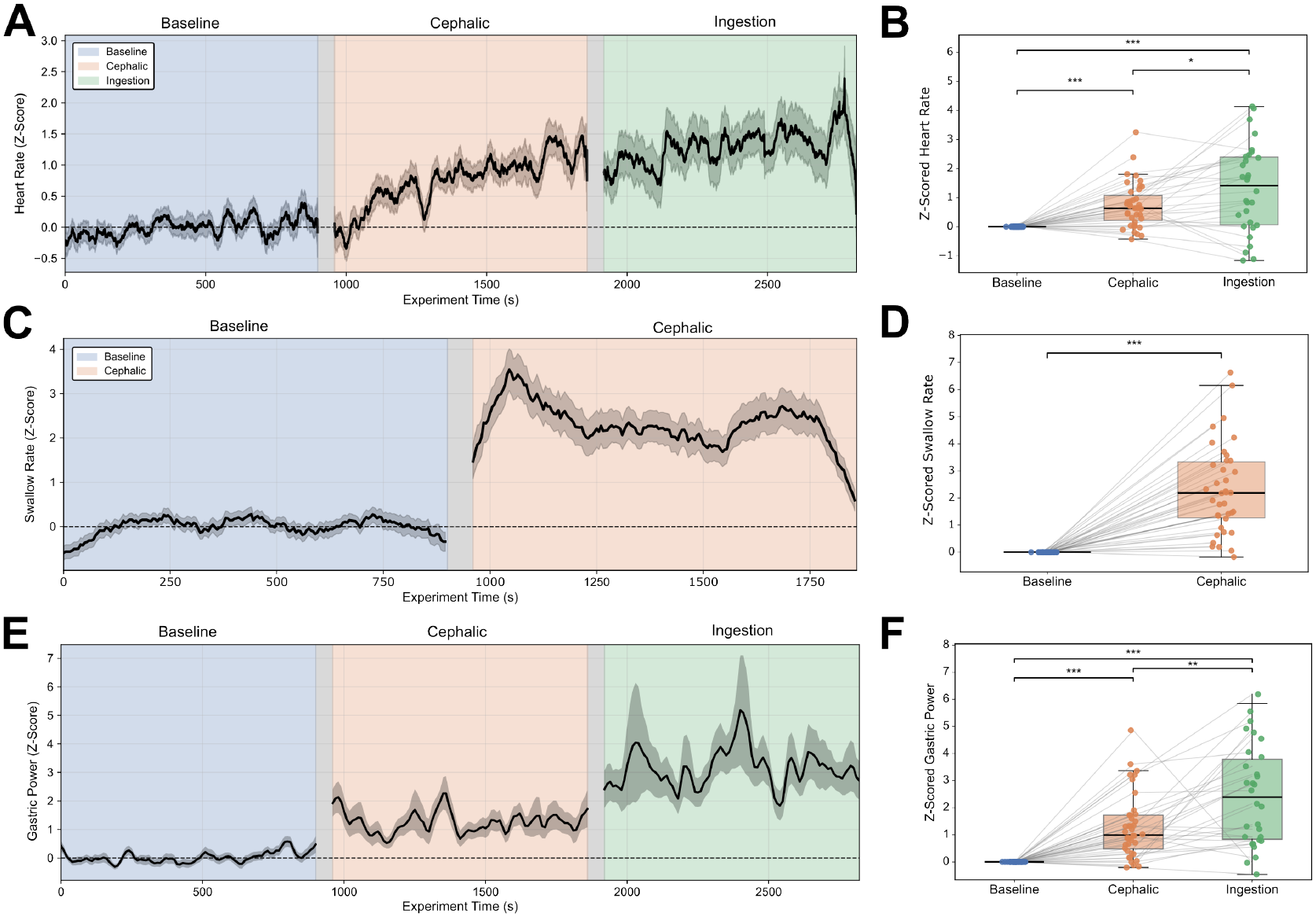
Visceral physiological responses during food anticipation and post-meal consumption. (A,B) Time-resolved and participant-level heart rate across baseline, cephalic, and ingestion phases. (C,D) Corresponding swallow rate during baseline and cephalic phases. (E,F) Time-resolved and participant-level gastric-band power (0.03–0.07 Hz) across all three phases. Temporal trajectories show the group mean *±* SEM, z-scored relative to baseline, with experimental phases indicated by blue, orange, and green shading, respectively. Boxplots display the median and interquartile range with paired participant observations. Primary baseline–cephalic phase comparisons used paired *t*-tests, while secondary ingestion phase comparisons were Holm-adjusted. *^∗^p <* 0.05, *^∗∗^p <* 0.01, *^∗∗∗^p <* 0.001.

Following nutrient ingestion, heart rate and gastric-band power increased further relative to the cephalic phase (Holm-adjusted *p* = 0.0185 and *p* = 0.0021, respectively) and remained significantly elevated relative to baseline (Holm-adjusted *p <* 0.001 for both). Repeated-measures ANOVAs confirmed significant overall effects of digestive phase on heart rate (*F* (2, 58) = 17.23, *p <* 0.001, partial *η*^2^ = 0.37) and gastric-band power (*n* = 32, *F* (2, 62) = 20.27, *p <* 0.001, partial *η*^2^ = 0.395). Together, these findings demonstrate that food anticipation elicits concurrent changes in swallow, cardiovascular, and gastric physiology, with distinct temporal trajectories extending into the ingestion phase. Whereas swallow activity increased rapidly in response to food cues, heart rate and gastric myoelectric power exhibited more sustained changes that persisted or increased further following nutrient intake.

### 3.2 The cephalic phase induces transient frequency-specific reorganization of cortical oscillatory activity

To characterize cortical responses during food anticipation and post-meal consumption, we quantified normalized whole-brain EEG spectral power across theta, alpha, and beta frequencies (Fig. 2). During the cephalic phase, theta power increased significantly relative to baseline (*n* = 40, *t*(39) = 3.90, *p <* 0.001, Cohen’s *d_z_* = 0.62; Fig. 2B), with frontocentral increases evident in the corresponding scalp topographies (Fig. 2A). Alpha power decreased significantly (*n* = 40, *t*(39) = *−*7.93, *p <* 0.001, Cohen’s *d_z_* = *−*1.25; Fig. 2D), accompanied by widespread alpha attenuation across the scalp (Fig. 2C). Beta power increased significantly (*n* = 40, *t*(39) = 4.45, *p <* 0.001, Cohen’s *d_z_* = 0.70; Fig. 2F), with bilateral frontocentral and left frontotemporal increases evident in the scalp topographies (Fig. 2E).

**Figure 2:**
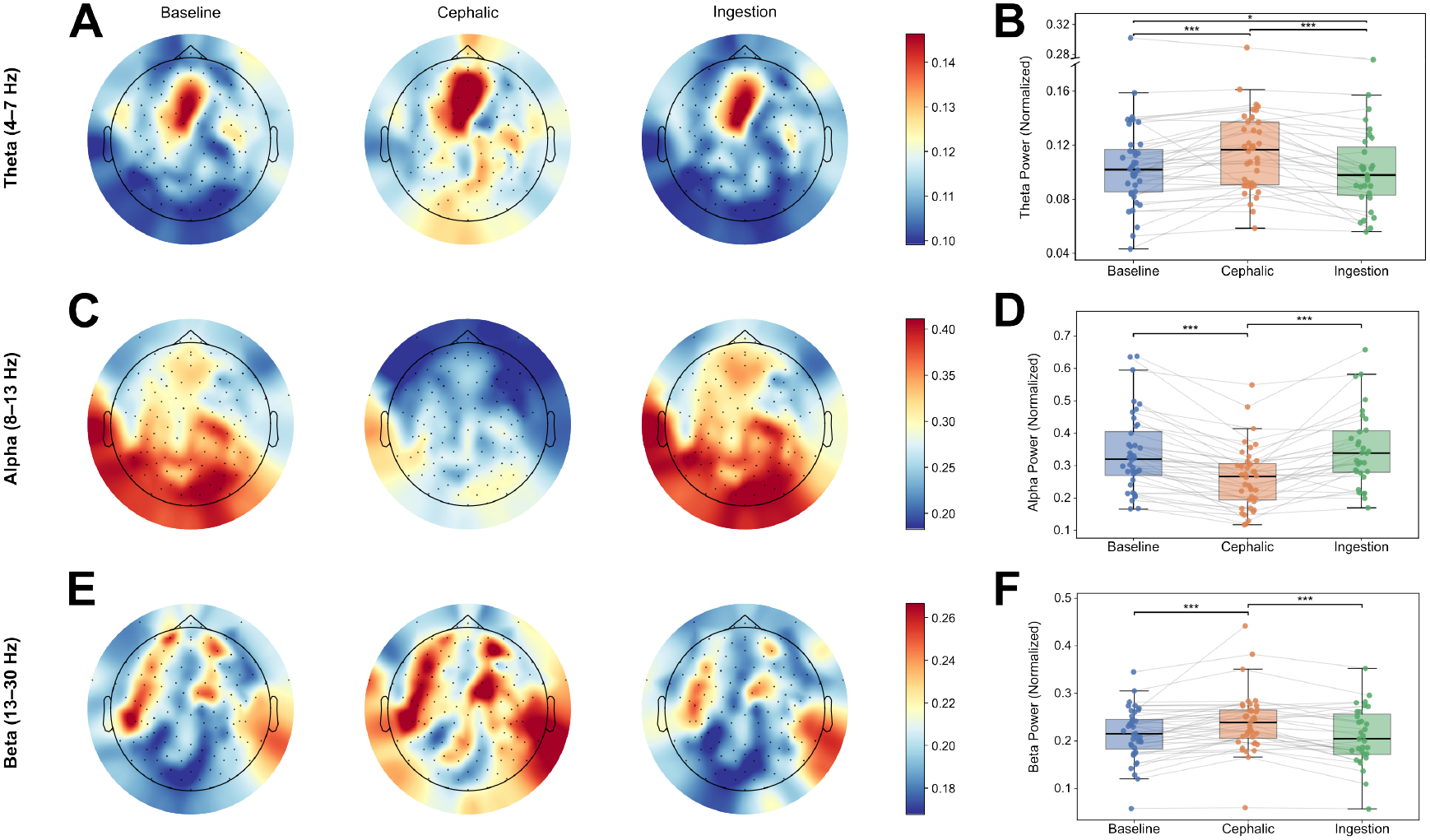
Frequency-specific changes in cortical oscillatory activity during food anticipation and following meal consumption. (A,C,E) Group-average scalp topographies of normalized theta (4– 7 Hz), alpha (8–13 Hz), and beta (13–30 Hz) power, respectively, across baseline, cephalic, and ingestion phases. (B,D,F) Corresponding participant-level mean normalized power across experimental phases, with paired observations overlaid. Boxplots show the median and interquartile range. Primary baseline–cephalic phase comparisons used paired *t*-tests, while secondary ingestion phase comparisons were Holm-adjusted. *^∗^p <* 0.05, *^∗∗^p <* 0.01, *^∗∗∗^p <* 0.001.

Following meal consumption, theta and beta power decreased significantly relative to the cephalic phase (Holm-adjusted *p <* 0.001 for both; Fig. 2B,F), while alpha power increased (Holm-adjusted *p <* 0.001; Fig. 2D). Alpha and beta power returned toward baseline, with no significant baseline-to-ingestion differences (alpha: Holm-adjusted *p* = 0.785; beta: Holm-adjusted *p* = 0.701), whereas theta power decreased below baseline (Holm-adjusted *p* = 0.0289). Repeated-measures ANOVAs confirmed significant overall effects of digestive phase across all three frequency bands (theta: *F* (2, 60) = 22.73, *p <* 0.001; alpha: *F* (2, 60) = 27.62, *p <* 0.001; beta: *F* (2, 60) = 12.66, *p <* 0.001). Together, these findings demonstrate a transient, frequency-specific cortical response to food anticipation that largely reverses following ingestion, with theta activity exhibiting a distinct post-ingestive reduction.

### 3.3 Gastric and cortical network synchrony decreases during the cephalic phase and recovers following ingestion

To characterize changes in within-system synchronization, we quantified phase-locking value (PLV) across gastric and cortical electrode pairs during the baseline, cephalic, and ingestion phases. Whole-network gastric synchrony was calculated by averaging PLV across the 28 unique pairs of eight EGG electrodes (Fig. 3A), using overlapping 150-s windows advanced in 30-s steps (Fig. 3B). Cortical synchrony was similarly assessed across scalp EEG electrode pairs within theta, alpha, and beta frequency bands (Fig. 4).

**Figure 3:**
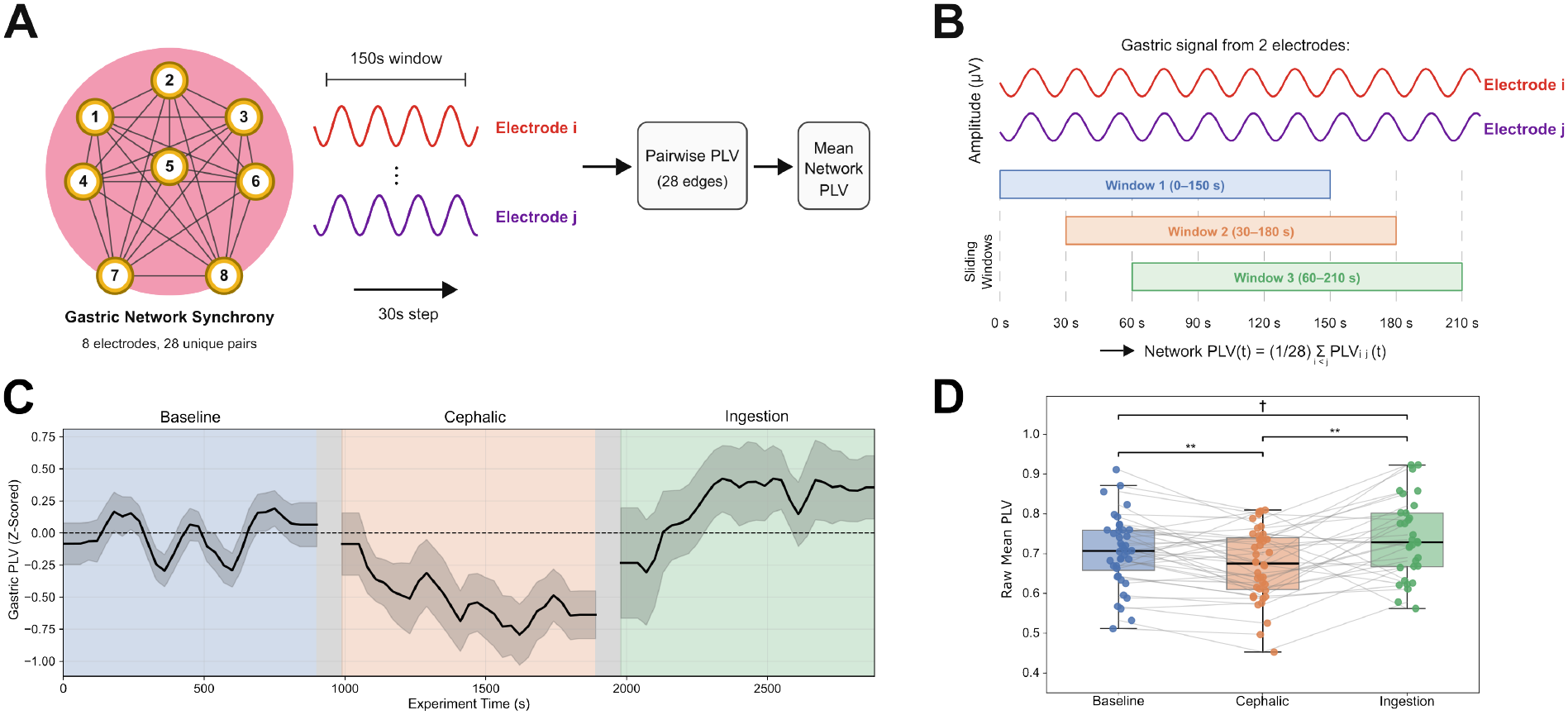
Dynamic reorganization of gastric network synchrony during food anticipation and post-meal consumption. (A) Gastric network synchrony was quantified as the mean phase-locking value (PLV) across 28 unique pairs of eight EGG electrodes. (B) PLV was calculated using overlapping 150-s windows advanced in 30-s steps between each pair of electrodes. (C) Time-resolved gastric PLV, z-scored to baseline, showing the group mean *±* SEM across experimental phases. (D) Participant-level mean raw gastric PLV, with paired observations overlaid. Boxplots show the median and interquartile range. Primary baseline–cephalic phase comparisons used paired *t*-tests, while secondary ingestion phase comparisons were Holm-adjusted. *^∗∗^p <* 0.01; *†* indicates a trend (0.05 *≤ p <* 0.10).

**Figure 4:**
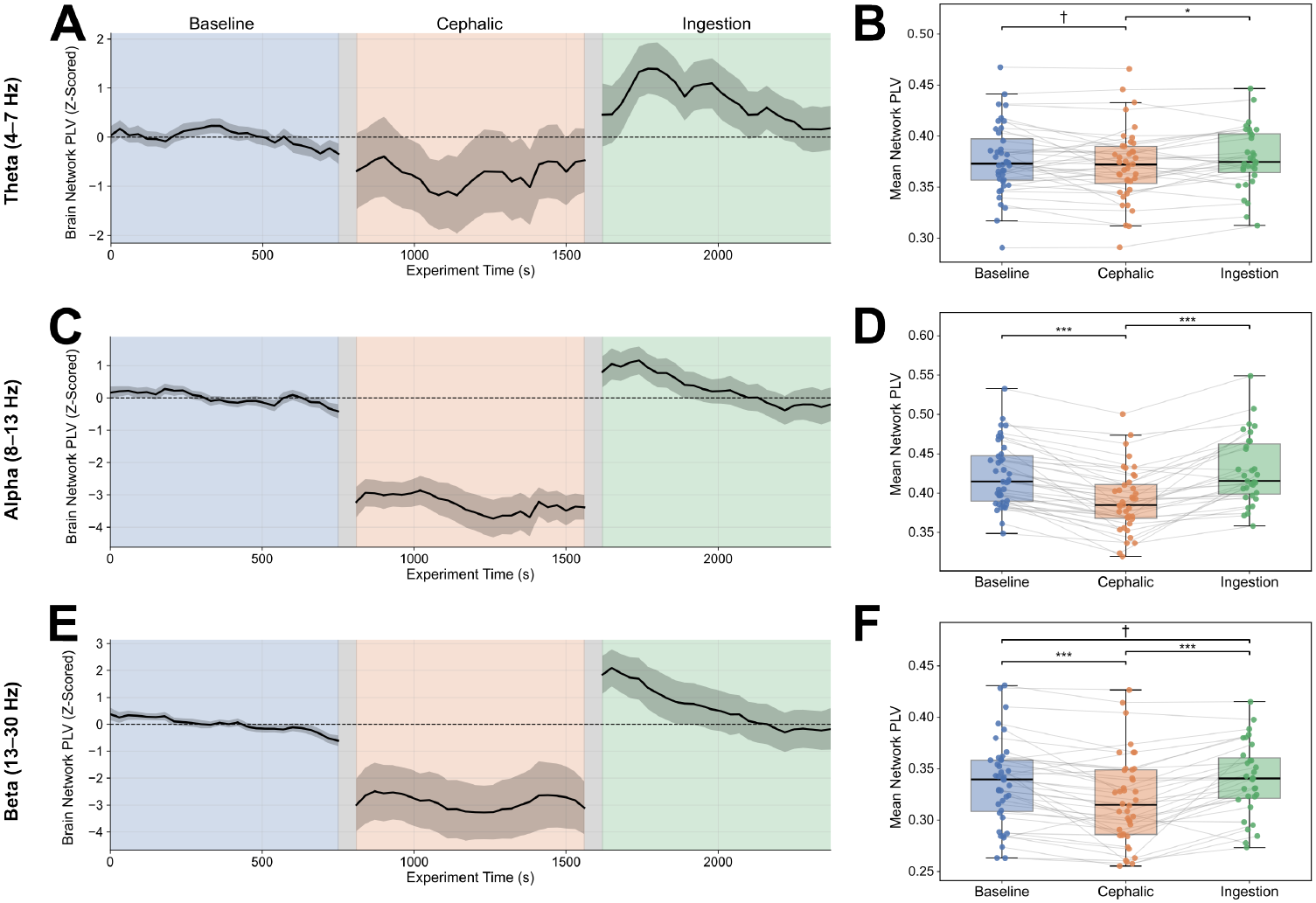
Cortical network synchrony decreases during food anticipation and recovers following ingestion. (A,C,E) Time-resolved whole-head phase-locking value (PLV) across theta (4–7 Hz), alpha (8–13 Hz), and beta (13–30 Hz) bands, respectively, z-scored to baseline. (B,D,F) Corresponding participant-level mean raw PLV across experimental phases. PLV was averaged across scalp EEG electrode pairs. Time series show the group mean *±* SEM; boxplots show the median and interquartile range with paired observations. Primary baseline–cephalic phase comparisons used paired *t*-tests, while secondary comparisons involving the ingestion phase were Holm-adjusted. *^∗^p <* 0.05, *^∗∗∗^p <* 0.001; *†* indicates a trend (0.05 *≤ p <* 0.10).

Gastric network synchrony progressively decreased during the cephalic phase and subsequently recovered during ingestion, as shown by the time-resolved PLV trajectory (Fig. 3C). Mean gastric PLV was significantly lower during anticipation than baseline (*n* = 40, *t*(39) = *−*3.07, *p* = 0.0039, Cohen’s *d_z_* = *−*0.485; Fig. 3D). Gastric PLV subsequently increased during ingestion relative to the cephalic phase (Holm-adjusted *p* = 0.0041), with a nonsignificant tendency toward greater synchrony than baseline (Holm-adjusted *p* = 0.0719; Fig. 3D). Repeated-measures ANOVA confirmed a significant overall effect of digestive phase (*F* (2, 62) = 7.61, *p* = 0.0011, partial *η*^2^ = 0.197).

Cortical network synchrony exhibited a parallel but frequency-dependent pattern. Time-resolved trajectories revealed reductions in theta-, alpha-, and beta-band PLV during anticipation, followed by increases during ingestion (Fig. 4A,C,E). Participant-level comparisons demonstrated significant reductions in mean alpha- and beta-band PLV relative to baseline (alpha: *n* = 40, *t*(39) = *−*8.52, *p <* 0.001, Cohen’s *d_z_* = *−*1.35; Fig. 4D; beta: *n* = 40, *t*(39) = *−*4.94, *p <* 0.001, Cohen’s *d_z_* = *−*0.78; Fig. 4F), whereas theta PLV showed a smaller, nonsignificant reduction (*t*(39) = *−*1.69, *p* = 0.099; Fig. 4B). During ingestion, synchrony increased significantly across all three bands relative to the cephalic phase (theta: Holm-adjusted *p* = 0.040; alpha and beta: Holm-adjusted *p <* 0.001; Fig. 4B,D,F), with a rapid increase at ingestion-phase onset followed by a gradual return toward baseline (Fig. 4A,C,E). Mean ingestion PLV did not significantly differ from baseline in any band (theta: Holm-adjusted *p* = 0.255; alpha: *p* = 0.267; beta: *p* = 0.074). Repeated-measures ANOVAs confirmed significant overall effects of digestive phase on theta (*F* (2, 60) = 3.57, *p* = 0.034, partial *η*^2^ = 0.106), alpha (*F* (2, 60) = 36.01, *p <* 0.001, partial *η*^2^ = 0.546), and beta synchrony (*F* (2, 60) = 24.12, *p <* 0.001, partial *η*^2^ = 0.446). Together, these findings reveal parallel reductions in gastric and cortical network synchrony during food anticipation, followed by rapid resynchronization with the onset of the ingestion phase.

### 3.4 Gastric–cortical phase-amplitude coupling increases during food anticipation

To determine whether food anticipation altered gastric–cortical interactions, we quantified phase–amplitude coupling (PAC) between gastric slow-wave phase and cortical oscillatory amplitude using mutual information (MI). We assessed mean MI and positive MI area under the curve (AUC) above each participant’s baseline mean to characterize average coupling strength and cumulative above-baseline coupling elevations, respectively, across theta, alpha, and beta frequencies. Regional analyses were performed across frontal, parietal, bilateral temporal, and occipital regions, with regional comparisons corrected using the Benjamini–Hochberg false discovery rate (FDR) procedure.

Beta gastric–cortical PAC increased during food anticipation, with time-resolved coupling reaching its highest values toward the end of the cephalic phase before declining following meal consumption (Fig. 5A). Both mean beta MI (*n* = 40, *p* = 0.0046, Cohen’s *d_z_ ≈* 0.47; Fig. 5C, left) and positive MI AUC above baseline (*p* = 0.0023, *d_z_ ≈* 0.51; Fig. 5C, right) increased significantly during the cephalic phase. Regional analyses revealed significant increases in mean beta MI across frontal, parietal, and bilateral temporal regions (ΔMI = +0.0029–0.0037, FDR-adjusted *p* = 0.011–0.015; Fig. 5E, left), whereas positive MI AUC increased significantly across all five regions (ΔAUC = +1.59–3.20, FDR-adjusted *p* = 0.0049–0.033; Fig. 5E, right). These findings indicate a broadly distributed anticipatory increase in beta gastric–cortical coupling.

**Figure 5:**
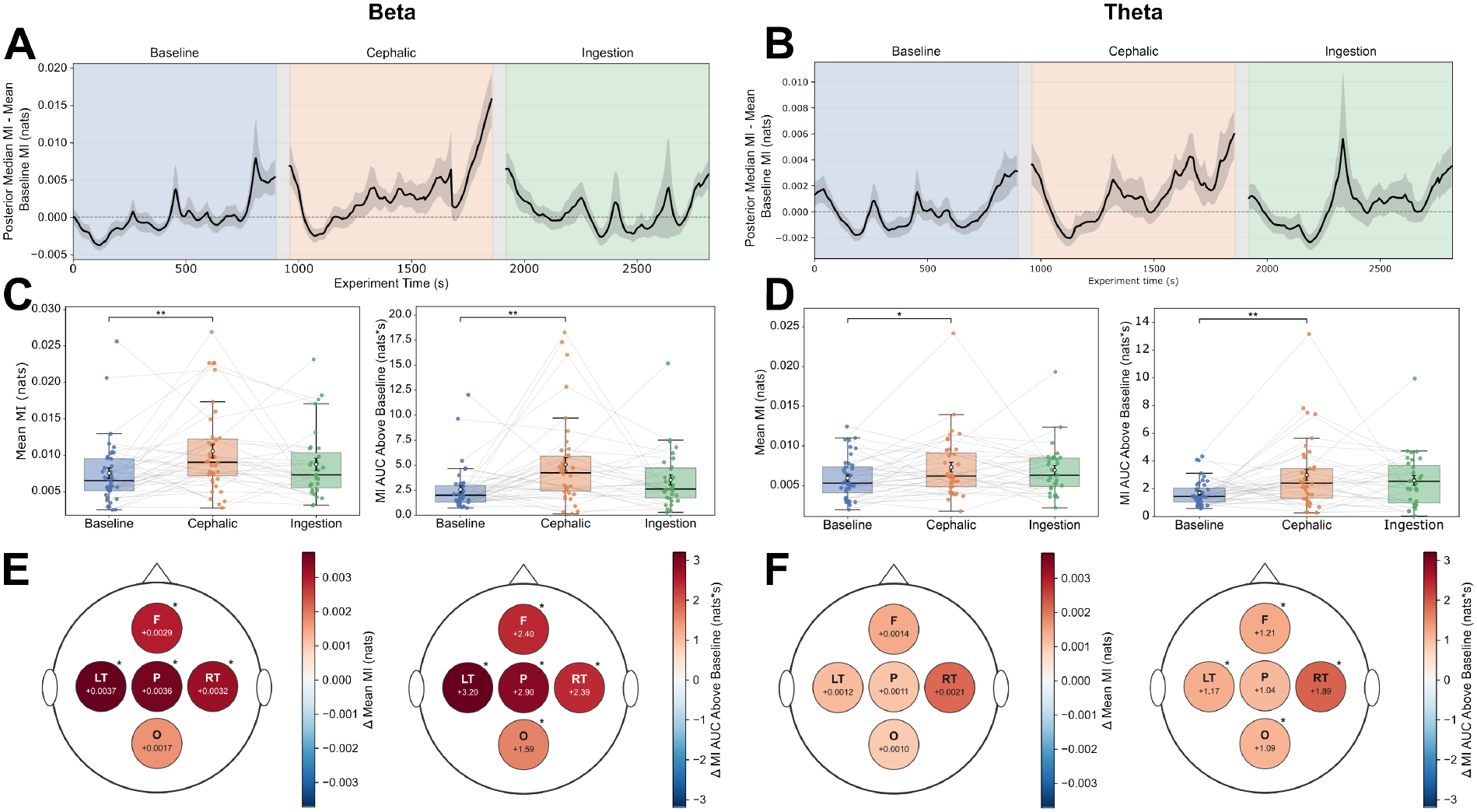
Gastric–cortical phase–amplitude coupling increases during food anticipation. (A,B) Time-resolved beta (13–30 Hz) and theta (4–7 Hz) PAC, respectively, quantified using posterior median mutual information (MI) after subtracting each participant’s baseline mean. (C,D) Participant-level mean MI (left) and positive excess MI area under the curve (AUC; right) for beta and theta, respectively, across experimental phases. (E,F) Regional cephalic-minus-baseline changes in mean MI (left) and positive excess AUC (right) across frontal, parietal, bilateral temporal, and occipital regions. Colors represent effect magnitudes, with asterisks indicating FDR-corrected significance. Time series show the group mean *±* SEM; boxplots show the median and interquartile range with paired observations. Primary baseline–cephalic phase comparisons used paired *t*-tests, with regional *p*-values corrected using the Benjamini–Hochberg false discovery rate procedure. *^∗^p <* 0.05, *^∗∗^p <* 0.01.

Theta PAC similarly increased during food anticipation, with time-resolved coupling remaining elevated into the ingestion phase (Fig. 5B). Both mean theta MI (*n* = 40, *p* = 0.0274, *d_z_ ≈* 0.36; Fig. 5D, left) and positive MI AUC above baseline (*p* = 0.0038, *d_z_ ≈* 0.49; Fig. 5D, right) increased significantly during the cephalic phase. Regional changes in mean theta MI were positive across all five regions (ΔMI = +0.0010–0.0021; Fig. 5F, left), although none survived FDR correction; the right temporal region exhibited a trend toward increased coupling (FDR-adjusted *p* = 0.055). In contrast, positive MI AUC increased significantly across frontal, bilateral temporal, and occipital regions (ΔAUC = +1.09–1.89, FDR-adjusted *p* = 0.019–0.020; Fig. 5F, right), indicating a more spatially distributed cumulative above-baseline response than was evident from changes in average coupling strength. These regional AUC increases remained significant across all 40 leave-one-participant-out sensitivity analyses, indicating that the effects were not driven by any single participant.

Alpha gastric–cortical PAC exhibited a comparatively weaker anticipatory response. Although mean alpha MI did not significantly change during the cephalic phase (*n* = 40, *p* = 0.400), whole-head positive MI AUC above baseline increased significantly (*p* = 0.023, Cohen’s *d_z_* = 0.37). Regional analyses revealed trends toward increased positive MI AUC across parietal, bilateral temporal, and occipital regions (FDR-adjusted *p* = 0.057), although none survived correction for multiple comparisons.

Together, these findings indicate frequency-dependent increases in gastric–cortical coupling during food anticipation, with the strongest regional effects observed in beta frequencies and more widespread cumulative above-baseline responses for theta frequencies.

### 3.5 Cephalic-phase gastric and cortical responses relate to subjective appetite

To examine the relationship between cephalic-phase physiology and subjective appetite, we assessed associations between gastric and cortical responses and participants’ self-reported hunger, satiety, and craving levels (*n* = 40). Subjective ratings revealed a significant increase in hunger following food anticipation (*t*(39) = 17.74, *p <* 0.001) and a decrease in satiety (*t*(39) = *−*8.50, *p <* 0.001; Fig. 6A). Greater cephalic-phase increases in gastric myoelectric power were associated with greater increases in hunger (*β* = 9.80, 95% CI [4.12, 15.48], *p <* 0.001; Fig. 6B), greater craving (*β* = 6.35, 95% CI [1.49, 11.20], *p* = 0.010; Fig. 6C), and larger reductions in satiety (*β* = *−*4.54, 95% CI [*−*8.84*, −*0.25], *p* = 0.038).

**Figure 6:**
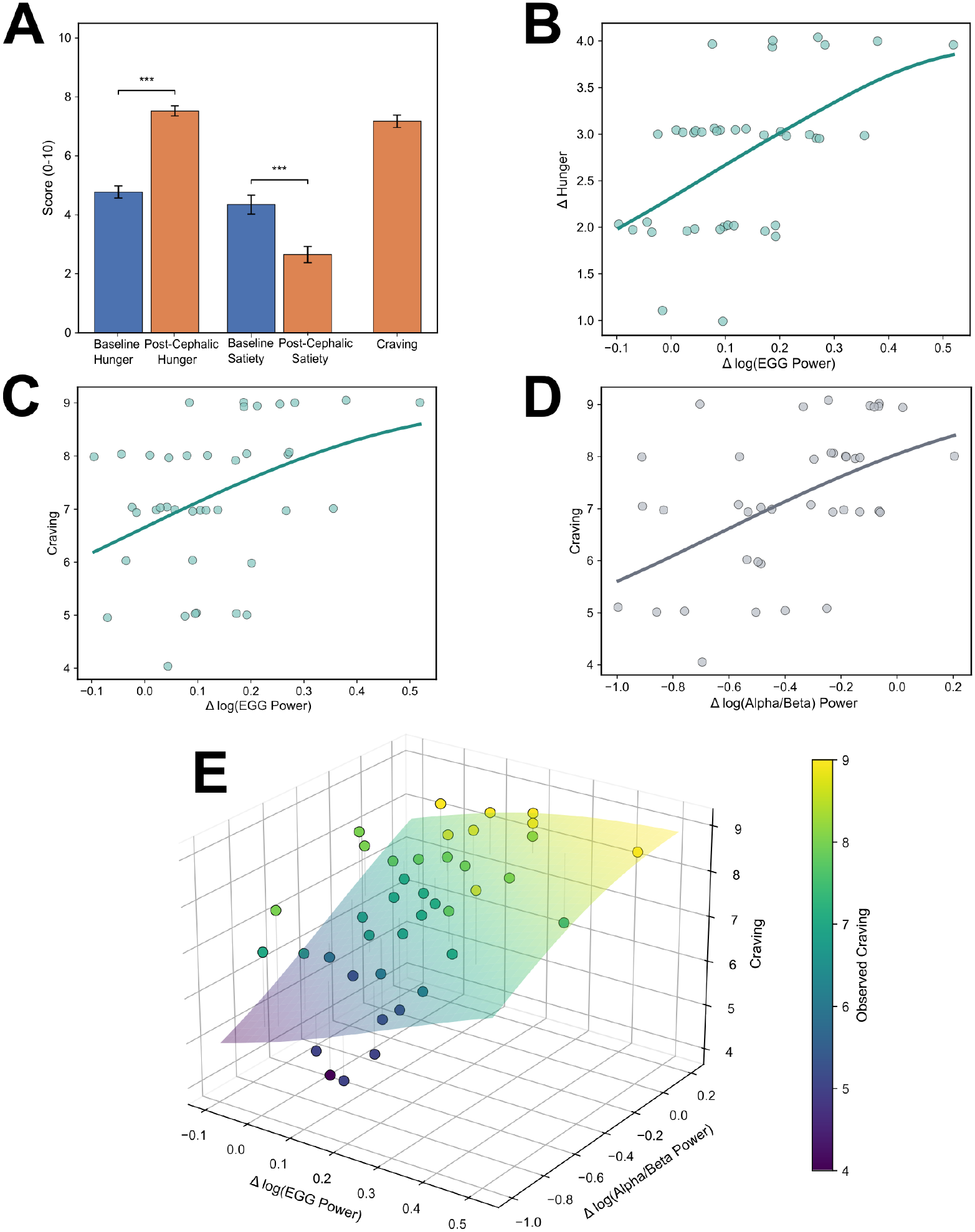
Cephalic-phase gastric and cortical responses relate to subjective appetite. (A) Self-reported hunger and satiety before and after food anticipation. (B,C) Associations of baseline-to-cephalic changes in gastric myoelectric power with changes in hunger (B) and post-cephalic craving ratings (C). (D) Association between changes in frontal log(alpha/beta) power ratio and craving. (E) Multivariable ordinal regression showing independent associations of gastric power and frontal log(alpha/beta) changes with craving. The frontal region is shown as a representative example; cortical associations remained significant across all five regions after FDR correction. Regional analyses used ordinal logistic regression with Benjamini–Hochberg FDR correction. Multivariable analyses included *n* = 40 participants. Error bars in (A) indicate SEM.

Cortical spectral changes were particularly associated with craving. Although alpha power decreased and beta power increased overall during anticipation, greater craving was associated with smaller alpha reductions and smaller beta increases across all five cortical regions after FDR correction (alpha: *p*_FDR_ = 0.0049–0.0081; beta: *p*_FDR_ = 0.0045–0.0235). Consistent with these findings, more positive cephalic-phase changes in the log(alpha/beta) power ratio were associated with greater craving in frontal (*β* = 3.69, *p*_FDR_ = 0.0070; Fig. 6D), right temporal, parietal, and occipital regions (*p*_FDR_ = 0.0184), whereas the left temporal association did not survive FDR correction (*p*_FDR_ = 0.0508). No regional associations between the cortical ratio and hunger or satiety survived FDR correction.

We next used multivariable ordinal regression to determine whether gastric and cortical responses were independently associated with participants’ self-reported craving. Models included cephalic-phase changes in gastric myoelectric power and cortical log(alpha/beta) ratio as simultaneous predictors, with separate models for each cortical region. Both predictors remained independently associated with craving across all five regional models. After accounting for cortical spectral changes, greater gastric power increases were associated with greater craving (adjusted OR = 2.16–2.57, *p* = 0.0070–0.024). Conversely, after accounting for gastric activity, more positive cortical log(alpha/beta) changes were associated with greater craving (adjusted OR = 2.08–2.90, *p*_FDR_ = 0.014–0.030), with the strongest cortical association observed in the frontal region (*OR* = 2.90, 95% CI [1.45, 5.81], *p*_FDR_ = 0.014; Fig. 6E).

Together, these findings demonstrate distinct but complementary relationships between physiological anticipation and subjective appetite. Gastric responses were associated with hunger, satiety, and craving, whereas cortical spectral responses were selectively related to craving. Importantly, gastric and cortical responses independently accounted for variation in craving, with the cortical association distributed across multiple regions and strongest over the frontal cortex. Collectively, our results suggest that individual differences in subjective appetite during food anticipation are associated with distinct yet complementary visceral and cortical components of the cephalic phase response.

## 4 Discussion

The cephalic phase is traditionally understood through the digestive responses that occur before food reaches the gastrointestinal tract. Here we revise this understanding by showing that food anticipation engages a coordinated physiological state spanning cortical, gastric, and autonomic systems. Food-related sensory cues increase swallow and heart rate as well as gastric myoelectric activity, while producing distinct frequency-specific changes in cortical activity. More strikingly, these responses are accompanied by opposing changes in physiological coordination: synchrony within gastric and cortical networks decreases, while coordination between the brain and stomach increases. These findings suggest that anticipation does not simply activate multiple physiological systems in parallel, but reorganizes how their activity is coordinated before nutrients enter the gastrointestinal tract. Moreover, gastric and cortical responses showed distinct relationships with hunger, satiety, and craving, indicating that this coordinated anticipatory state encompasses varying aspects of appetite. Taken together, our results reveal a broad organization of the human cephalic response in which sensory anticipation is accompanied by coordinated changes in central and peripheral physiology before ingestion begins.

The physiological changes observed during food anticipation are consistent with the established role of cholinergic vagal pathways in cephalic digestive preparation. Food-related sensory cues engage forebrain and brainstem autonomic circuits, including the DMV, which provides cholinergic efferent signaling to the gastrointestinal tract [8, 3, 16]. Although previous studies have established the importance of these pathways in anticipatory digestive and autonomic responses, our findings extend this work by simultaneously characterizing gastric electrical activity, cortical oscillations, and cardiovascular responses during food anticipation. This multimodal approach provides a broader view of the physiological changes accompanying cephalic digestive preparation.

A particularly notable finding is the increase in gastric electrical activity during food anticipation, which persisted following ingestion. Gastric-band power increased during the cephalic phase, consistent with previous sham-feeding studies demonstrating vagally mediated increases in the gastric 3-cpm rhythm [17]. The gastric slow wave, generated by interstitial cells of Cajal, provides the electrical timing framework for gastric motor activity and is modulated by enteric and autonomic pathways [18, 19]. The anticipatory increase in gastric power likely reflects modulation of gastric electrical activity by the cholinergic pathways recruited during the cephalic phase.

Alongside these peripheral responses, food anticipation induced a distinct cortical oscillatory state characterized by increased theta and beta power and widespread alpha attenuation. Frontocentral theta enhancement may reflect olfactory and attentional processing, while alpha attenuation is consistent with increased processing of behaviorally relevant sensory information [20, 21, 22, 23]. Beta enhancement may additionally reflect top-down sensorimotor processes involved in preparation for consumption [24]. Together, these frequency-specific changes suggest the recruitment of distributed cortical processes supporting sensory processing, attention, motivation, and motor preparation during food anticipation [14, 13, 25]. Notably, cortical spectral activity returned toward baseline following ingestion, whereas gastric power remained elevated. This divergence suggests that the cortical response is particularly sensitive to the anticipatory period, while gastric activation extends into post-ingestive processing. The transition from anticipation to ingestion therefore involves not simply a change in response magnitude, but a shift in the relative engagement of central and peripheral physiological systems.

A particularly notable finding was the reduction in synchronization within gastric and cortical networks during food anticipation. Phase synchrony progressively decreased across gastric recording sites, accompanied by pronounced reductions in cortical alpha- and beta-band synchrony and a smaller reduction in theta synchrony. Following ingestion, synchronization increased rapidly within both systems. These findings demonstrate that increased physiological activity during anticipation does not necessarily correspond to increased within-system synchronization, suggesting distinct patterns of physiological organization during anticipation and ingestion.

In the cortex, reduced large-scale synchronization may reflect a shift toward more flexible and differentiated neural processing, allowing distributed neuronal populations to respond to sensory demands without remaining locked into a common oscillatory pattern [26, 27]. During food anticipation, this flexibility may facilitate the integration of visual and olfactory cues with attentional, motivational, and motor-related processes. Importantly, reduced phase synchronization does not necessarily imply disorganized neural activity, as functional coordination may be maintained through other forms of interaction [28].

The parallel gastric response raises the possibility that anticipation also changes the organization of peripheral electrical activity. Similar to cortical desynchronization, reduced gastric synchrony may reflect a shift toward a more adaptable physiological state during food anticipation. Gastric slow waves normally propagate across the stomach to provide the electrical framework for coordinated motor activity [18, 29]. Thus, desynchronization across abdominal recordings could reflect changes in the spatial or temporal organization of gastric electrical activity, potentially facilitating responsiveness to anticipatory signaling as the stomach prepares for incoming food. The rapid increase in gastric synchrony following ingestion may reflect a transition toward more coordinated electrical activity as the stomach shifts from anticipating food to processing it. Together, the parallel changes in gastric and cortical synchronization suggest that food anticipation and ingestion engage distinct patterns of physiological organization across central and peripheral systems.

Crucially, reduced synchronization within individual systems is accompanied by increased coordination between them. Phase–amplitude coupling revealed frequency- and spatially dependent changes, with particularly prominent anticipatory effects involving theta and beta activity. Thus, while gastric and cortical networks become less internally synchronized, their activity becomes more closely related across the gut– brain axis. This dissociation suggests that food anticipation does not involve a uniform increase or decrease in physiological coordination, but rather a redistribution of coordination across organizational scales. Before nutrients arrive, the brain must process food-related sensory information and organize preparatory responses while the gastrointestinal system responds to the expectation of food. The coexistence of reduced within-system synchrony and enhanced cross-system coupling may allow individual systems to retain flexibility in their local activity while remaining coordinated across the brain–body axis. This interpretation is consistent with evidence that reduced synchronization within networks can coexist with structured coordination at other spatial or functional scales [28].

The frequency specificity of gastric–cortical coupling further suggests that these interactions relate to different components of anticipatory processing. Previous studies have emphasized gastric–cortical coordination during resting states, particularly involving alpha-band activity [30, 31, 32]. In contrast, the prominent theta- and beta-related coupling observed during anticipation may reflect additional engagement of olfactory, attentional, and sensorimotor processes associated with impending food consumption. These findings suggest that gastric–cortical coupling is dynamically modulated by behavioral state, extending beyond the resting conditions in which it has primarily been studied.

Although food anticipation elicited coordinated physiological changes in all participants, the magnitude of these responses varied between individuals and was related to differences in hunger, satiety, and craving. Importantly, gastric and cortical responses were not associated with these measures in the same way. Greater anticipatory increases in gastric myoelectric power were associated with greater hunger and craving and lower satiety, whereas cortical spectral changes were more selectively related to craving. This distinction suggests that the physiological response to anticipated food contains multiple components that provide different information about appetite.

The broader association between gastric activity and appetite is consistent with the stomach’s role in digestive preparation and homeostatic regulation. Previous work has demonstrated that cephalic gastric responses vary with the appetitive properties of food and can depend on vagal signaling [17, 33, 34]. We now show that individual differences in anticipatory gastric electrical activity are related to hunger, satiety, and craving. Rather than representing a uniform response to sensory cues, gastric recruitment during anticipation therefore varies with the individual’s appetite, suggesting that anticipatory gastric electrophysiology carries information relevant to appetitive state.

Cortical responses, in contrast, showed a more selective relationship with craving. Interestingly, although food anticipation produced a group-level decrease in alpha power and increase in beta power, participants reporting greater craving exhibited smaller changes in both frequencies. Greater craving was therefore associated with a less pronounced shift from alpha toward beta activity, particularly over frontal cortical regions. This finding is notable because it suggests that stronger craving does not necessarily correspond to a larger cortical response to food-related cues. Instead, the magnitude of cortical spectral reorganization likely depends on how individuals evaluate and respond to anticipated food.

The frontal predominance of these associations is consistent with the involvement of prefrontal and meso-corticolimbic circuits in food valuation, motivated attention, and craving [35, 36, 37, 38, 39]. Dopaminergic signaling within these circuits contributes to incentive motivation and the attribution of motivational significance to anticipated rewards, providing a possible framework for understanding individual differences in craving. In contrast, cholinergic vagal pathways mediate peripheral digestive preparation during the cephalic phase. The divergent cortical and gastric associations likely reflect partially distinct motivational and autonomic processes recruited during anticipation, rather than opposing dopaminergic and cholinergic activity. Although the physiological basis of the smaller cortical spectral changes associated with greater craving remains uncertain, our findings show that motivational state and the magnitude of cortical reorganization are not necessarily directly proportional.

Importantly, the gastric and cortical associations with craving are distinct. When gastric myoelectric power and regional cortical spectral measures were included simultaneously in multivariable models, both remained associated with craving. This indicates that the two signals provide complementary, rather than redundant, information about differences in craving between individuals. Gastric activity seems more closely related to the visceral and homeostatic aspects of appetite, whereas cortical oscillatory changes may reflect processes involved in sensory evaluation, motivation, and anticipated reward. These interpretations are not mutually exclusive, and the findings do not establish separate causal pathways. Rather, they suggest that craving is associated with distinguishable components of anticipatory brain–body physiology.

This distinction adds an important new dimension to our understanding of the cephalic phase. Although cortical and gastric systems become more closely coordinated during anticipation, their physiological responses are not interchangeable, nor do they show identical relationships with appetite. The same anticipatory state can therefore involve increased functional integration across systems while preserving distinct associations between each system and the individual’s experience of food anticipation. Understanding appetite will consequently require examining both central and visceral physiology, rather than relying on either alone.

Our findings have important implications for the study of vagally mediated gut–brain function in humans. Despite the central role of cholinergic vagal signaling in digestive regulation, descending vagal efferent activity is difficult to characterize non-invasively [40, 41]. Classical cephalic-phase studies have addressed this challenge through peripheral measures such as gastric secretion, insulin release, and pancreatic polypeptide responses, with vagotomy and cholinergic blockade demonstrating the importance of vagal pathways for several of these effects [11, 34, 42]. Although informative, these approaches generally characterize individual digestive outputs rather than the broader physiological organization accompanying anticipatory vagal control.

Our data suggest that simultaneous cortical, gastric, cardiac, and swallow measurements provide a more comprehensive way to characterize the human cephalic response. Rather than relying on a single peripheral endpoint, this approach captures several complementary features of food anticipation: changes in cortical processing, gastric electrical activity, autonomic and behavioral output, synchronization within physiological systems, and coordination between the brain and stomach. The distinct relationships of gastric and cortical responses with appetite further suggest that these signals capture different dimensions of anticipatory physiology. Together, they provide a potential multimodal signature of the physiological processes recruited before nutrient ingestion.

Such an approach may be particularly valuable in conditions involving altered autonomic regulation or gut–brain communication. Parkinson’s disease represents one important application because gastrointestinal dysfunction and autonomic abnormalities are common features of the disease, and pathological involvement of the DMV and enteric nervous system affect the pathways associated with cephalic digestive regulation [43]. Characterizing anticipatory physiological responses in Parkinson’s disease can help determine whether abnormalities in gut–brain function are reflected in altered gastric recruitment, cortical responses, or coordination between them. Similar approaches may eventually inform research into gastrointestinal dysmotility, metabolic disorders, and interventions modulating appetite or gut–brain signaling, including GLP-1 receptor agonists and vagal neuromodulation [44, 4, 5].

Several limitations are important when considering these applications. First, our measurements provide indirect indices of physiology associated with vagal regulation; they do not directly measure cholinergic efferent activity or establish the vagal specificity of the responses. Second, changes in phase locking and gastric–cortical coupling describe statistical relationships between recorded signals but cannot determine causal interactions, neural directionality, or the underlying physiological mechanisms. In particular, surface EGG synchronization cannot independently establish changes in gastric slow-wave propagation or motor function. Third, our observations in healthy adults do not establish the reliability or diagnostic utility of these measures in clinical populations. Future studies incorporating pharmacological or neuromodulatory interventions, repeated measurements, and comparisons with established autonomic and gastrointestinal assessments will be necessary to determine the mechanisms, reproducibility, and clinical relevance of this multimodal response.

More broadly, we suggest that anticipation itself provides an informative window into the organization of brain–body physiology. Before eating starts, food-related sensory cues are accompanied by changes in cortical, gastric, and autonomic activity, together with a reorganization of coordination within and between the brain and stomach. The cephalic phase therefore offers an opportunity to investigate how the nervous system and gastrointestinal tract become functionally engaged in expectation of a future physiological demand. By revealing these coordinated changes, our study provides a foundation for examining how anticipatory gut–brain organization is altered in disease and whether it can be modified through interventions targeting autonomic and gastrointestinal function.

## 5 Methods

In this study, we recruited 40 healthy participants with no diagnosed gastrointestinal, neurological, or psychiatric disorders. Participants were recruited from the local university and Stanford University campus using fliers and word of mouth. All participants provided written informed consent for the study protocol (no. 77495) approved by the Stanford Institutional Review Board (IRB).

### 5.1 Demographics

Demographic and dietary information were collected via self-report questionnaires in the days prior to the experimental session. Variables obtained included age, sex, date of birth, cultural background, typical meal timing patterns, preferred foods and cuisines, dietary restrictions, food allergies, and familiarity with a variety of food items. Questionnaire responses were used to guide individualized meal selection and ensure that food stimuli presented during the cephalic phase were familiar, culturally relevant, and highly appetitive for each participant.

### 5.2 Study Design

Participants completed a single 2-hour experimental study visit conducted at the Koret Neuroscience Laboratory at the Wu Tsai Neurosciences Institute. Experiments were conducted in an enclosed Faraday cage room within the laboratory to minimize electromagnetic interference. All participants arrived in a fasted condition following a 10-14-hour overnight fast. A schematic of the multimodal recording setup is shown in Figure 7A. Simultaneous electroencephalography (EEG), electrogastrography (EGG), electrocardiography (ECG), and swallow electromyography (EMG) recordings were acquired during baseline, cephalic, and ingestion phases to capture cortical, autonomic, and gastrointestinal responses to food anticipation and consumption.

**Figure 7:**
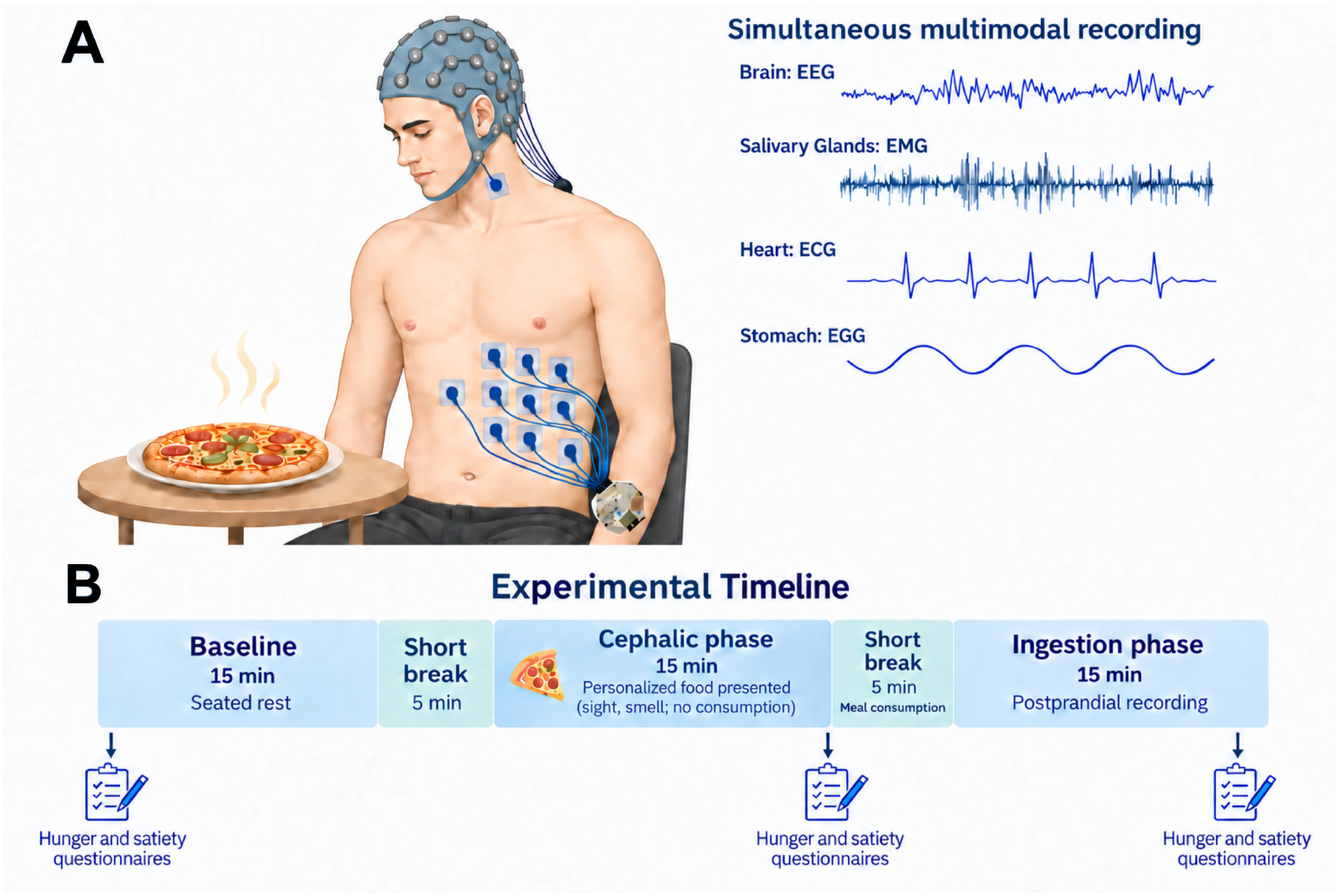
Multimodal recording setup and experimental timeline. (A) Simultaneous acquisition of cortical, autonomic, and gastrointestinal physiology during the experimental protocol, including electroen-cephalography (EEG), electrogastrography (EGG), electrocardiography (ECG), and swallow electromyography (EMG). (B) Following an overnight fast, participants completed 15-minute baseline and cephalic recordings, consumed their personalized meal during a brief break, and subsequently completed a 15-minute post-ingestive recording period (termed the ingestion phase).

The experimental timeline is illustrated in Figure 7B. The protocol consisted of three consecutive 15-minute periods, termed the baseline, cephalic, and ingestion phases, separated by brief 5-minute breaks. Participants first completed a 15-minute baseline recording while seated quietly at rest. During the cephalic phase, participants were presented with a personalized food stimulus and exposed to both visual and olfactory food cues to elicit anticipatory digestive responses. During the second break, participants consumed the presented meal. The ingestion phase was defined as the 15-minute post-ingestive recording period beginning after meal consumption, during which participants remained seated quietly at rest. Subjective appetite assessments were administered before baseline, immediately following the cephalic phase, and after completion of the ingestion phase.

After completion of setup on the day of testing, participants completed a brief questionnaire assessing demographic and clinical characteristics, including height, weight, sleep duration, medications taken, and acute gastrointestinal symptoms. Participants also reported the duration of their fasting period prior to testing. Subjective measures were assessed throughout the experiment. Hunger, satiety, stress, anxiety, and alertness were rated on a 0–10 scale (0, lowest; 10, highest) prior to baseline and following the cephalic and ingestion phases, while craving for the personalized meal was assessed following the cephalic phase. Participants also rated the appetizingness of the meal and reported how frequently they typically consumed it. Acute gastrointestinal symptoms initiated during each experimental phase were reported.

### 5.3 Multimodal Physiological Acquisition

During the study visit, EEG was recorded using a 128-channel high-density HydroCel Geodesic Sensor Net (GSN-HydroCel-128; Electrical Geodesics, Inc., Eugene, OR, USA) arranged according to the international 10–20 electrode placement system. Prior to cap placement, head circumference was measured using a flexible tape measure to determine the appropriate sensor net size. One of four adult cap sizes (47–51 cm, 51–54 cm, 55–58 cm, or 58–61 cm head circumference) was selected to ensure proper electrode positioning and fit. Electrode impedances were optimized using an electrolyte solution consisting of water, potassium chloride, and baby shampoo and were assessed prior to the start of the recording session and again immediately before the ingestion phase. To preserve electrode contact, participants wore a plastic shower cap over the sensor net throughout the recording period.

The Geodesic recording system was additionally interfaced with a 16-channel peripheral EMG amplifier, enabling synchronized acquisition of electrophysiological signals alongside the EEG. The peripheral unit supported 16 bipolar recording channels; here, two bipolar electrode pairs were used to measure submental swallow-related muscle activity. Swallow EMG was recorded using two surface electrodes placed over the left and right submental musculature, including the anterior digastric region. Each recording electrode was referenced to a corresponding electrode positioned over the ipsilateral mastoid process.

EGG was synchronously recorded using an OpenBCI electrophysiological recording system interfaced with 3M Red Dot Ag/AgCl electrodes, as used in previous EGG studies [45, 44, **?**, 32]. The configuration comprised eight recording electrodes, one reference electrode, and one bias electrode. Electrode placement was based on the abdominal montage described by Wolpert et al. [**?**], with a minor adjustment to the reference electrode position; the reference electrode was placed slightly inferior and lateral to the originally described location to avoid placement over the rib cage and improve electrode contact.

### 5.4 Signal Quality and Data Inclusion

Signal quality was evaluated independently for each physiological modality and experimental phase. Participants were excluded from analyses of a given modality when the baseline recording was of insufficient quality or when substantial artifact or signal loss compromised the recording throughout the experimental session. These exclusions were modality-specific and did not result in removal of the participant from analyses of other physiological measures with adequate signal quality. When poor signal quality or substantial artifact was restricted to the cephalic or ingestion phase, only the affected phase was excluded, while usable data from the remaining phases were retained. Consequently, sample sizes varied across modalities and experimental phases according to data availability and signal quality.

### 5.5 EEG Processing

Data were acquired at 500 Hz sampling frequency at 24-bit resolution. Event markers were recorded during the recording. The mff file retrieved from the NetStation app was analyzed in MATLAB using the EEGLAB toolbox. For each participant, continuous EEG recordings were segmented into the three experimental phases of interest (baseline, cephalic, and ingestion) prior to preprocessing. All preprocessing steps were performed separately for each phase in EEGLAB. Data were downsampled from 500 Hz to 250 Hz to reduce computational load while preserving the frequency content of interest. Channels exhibiting excessive noise, poor contact, or signal dropout were identified through visual inspection and marked as bad channels. The EEG data were then notch filtered between 55 and 65 Hz to attenuate line noise and bandpass filtered between 1 and 40 Hz using Hamming-windowed sinc finite impulse response (FIR) filters. Following filtering, the data were average referenced. Independent component analysis (ICA) was subsequently performed to identify and remove components associated with ocular, muscular, cardiac, and other non-neural artifacts. Post-ICA, previously identified bad channels were reconstructed using spherical spline interpolation. A final average reference was then applied to the interpolated dataset prior to subsequent postprocessing analyses.

#### 5.5.1 Extraction of EEG Band Power

Analyses focused on theta (4–7 Hz), alpha (8–13 Hz), and beta (13–30 Hz) activity, selected based on their established involvement in cortical responses to food-related cues and their relevance to attentional and sensorimotor processing. Spectral power within these frequency bands was quantified using Welch’s power spectral density (PSD) method. For each EEG channel, band-specific power was calculated by integrating the PSD across the corresponding frequency range, while total power was calculated by integrating the PSD from 1–30 Hz. Normalized band power was computed as the ratio of band-specific power to total 1–30 Hz power, thereby expressing oscillatory activity relative to overall broadband spectral power.

To characterize the spatial distribution of cortical spectral activity, phase-level scalp topographies were first generated for each frequency band. For each participant and experimental phase (baseline, cephalic, and ingestion), Welch PSD estimates were computed independently at each of the 128 scalp electrodes using 2-s segments. Relative theta, alpha, and beta power was calculated at each electrode as described above, yielding a participant-level spatial power map for each phase. Electrode-wise values were subsequently averaged across participants to generate group-level scalp topographies using the HydroCel 128-channel electrode montage. Difference topographies were additionally calculated to visualize phase-related changes in the spatial distribution of spectral power.

To characterize the temporal evolution of cortical spectral activity, normalized band power was additionally quantified within overlapping 60-s windows advanced in 30-s steps throughout each experimental phase. Within each window, relative theta, alpha, and beta power was calculated separately for each valid EEG electrode and then averaged across electrodes to obtain a single whole-head power estimate for each participant. The resulting whole-head power trajectories were aligned across experimental phases for group-level visualization.

For participant-level statistical analyses, whole-head relative power was summarized for each subject and phase by averaging the valid 60-s window-level whole-head power estimates across the experimental phase. Phase-related differences were evaluated according to the primary and secondary comparison framework described in the Statistical Analysis section.

#### 5.5.2 EEG Phase Synchronization

Whole-head cortical phase synchronization was quantified using phase locking value (PLV) within the same theta (4–7 Hz), alpha (8–13 Hz), and beta (13–30 Hz) frequency bands examined in the spectral power analyses. For each frequency band, preprocessed EEG signals were bandpass filtered using a fourth-order zero-phase Butterworth filter. Instantaneous phase was obtained independently for each EEG electrode using the Hilbert transform applied to the bandpass-filtered signal. Phase was estimated across the complete experimental phase prior to windowing to reduce Hilbert-transform edge effects within individual analysis windows. Whole-head phase synchronization was then quantified within overlapping 150-s windows advanced in 30-s steps throughout the baseline, cephalic, and ingestion phases. Within each window, PLV was calculated for every unique pair of scalp electrodes as the magnitude of the mean complex phase difference. Pairwise PLV values were subsequently averaged across the upper triangle of the electrode-by-electrode PLV matrix to obtain a single whole-head synchronization value for each analysis window.

For visualization of time-resolved changes, PLV values from each experimental phase were z-scored relative to each participant’s baseline by subtracting the mean PLV across the entire baseline phase and dividing by its standard deviation. The resulting trajectories were aligned across experimental phases for group-level visualization.

For participant-level statistical analyses, raw, non-normalized PLV values were averaged across all valid windows within each experimental phase to obtain one mean whole-head PLV value per participant, phase, and frequency band. Phase-related differences were evaluated according to the primary and secondary comparison framework described in the Statistical Analysis section.

### 5.6 EGG Processing

EGG signals were acquired using the OpenBCI graphical user interface (GUI), which received data wirelessly from the OpenBCI recording system via the USB radio dongle connected to the acquisition computer. The EGG signal was sampled at 250 Hz. To isolate gastric slow-wave activity, EGG signals were bandpass filtered between 0.03 and 0.07 Hz using a fourth-order zero-phase Butterworth filter, encompassing the gastric slow-wave frequency of approximately 0.05 Hz. Further artifact rejection was achieved by using a Wiener filter with window size equivalent to the inverse of the dominant frequency of each participant (approximately 20 s) as described previously [45]. To verify signal quality and identify the optimal gastric electrode to use for further analysis, PSD estimates were computed for each of the eight EGG electrodes during the baseline period using Welch’s method with a Hann window, 600-second segments, and 400-second overlap. Participants were included only if a dominant spectral peak was observed within 0.03–0.07 Hz. For subsequent analyses, gastric-band power was estimated across all eight EGG channels, and the channel exhibiting the median gastric-band power was selected as the representative gastric recording channel for each participant.

#### 5.6.1 Extraction of Gastric Band Power

Time-resolved gastric power from the selected channel was estimated using a sliding-window multitaper spectral approach. Power spectral density (PSD) estimates were computed in 250-s windows advanced in 10-s steps, and gastric-band power was quantified within the 0.03–0.07 Hz frequency range. Gastric power values were z-scored relative to each participant’s baseline period by subtracting the mean gastric power across the entire baseline phase and dividing by the corresponding baseline standard deviation. The resulting trajectories were aligned across experimental phases for group-level visualization. For participant-level statistical analyses, z-scored gastric power values were averaged across all valid windows within each experimental phase to obtain one mean gastric power value per participant and phase. Phase-related differences were evaluated according to the primary and secondary comparison framework described in the Statistical Analysis section.

#### 5.6.2 Gastric Phase Synchronization

Gastric network synchrony was quantified using PLV across all eight EGG electrodes within the gastric frequency band (0.03–0.07 Hz). For each participant, EGG data were segmented according to the baseline, cephalic, and ingestion phases and analyzed in overlapping 150-s windows advanced in 30-s steps. Within each window, EGG signals were downsampled from 250 Hz to 1 Hz, bandpass filtered from 0.03–0.07 Hz using a fourth-order zero-phase Butterworth filter, and instantaneous phase was extracted using the Hilbert transform. Pairwise PLV was calculated between all unique electrode pairs as the magnitude of the mean complex phase difference, and these values were averaged across the upper triangle of the electrode-by-electrode PLV matrix to obtain a single gastric network synchronization value for each analysis window.

For visualization of time-resolved changes, PLV values from each experimental phase were z-scored relative to each participant’s baseline period by subtracting the mean PLV across the entire baseline phase and dividing by the corresponding baseline standard deviation. The resulting trajectories were aligned across the baseline, cephalic, and ingestion phases for group-level visualization.

For participant-level statistical analyses, raw, non-z-scored PLV values were averaged across all valid windows within each experimental phase to obtain one mean gastric PLV value per participant and phase. Phase-related differences were evaluated according to the primary and secondary comparison framework described in the Statistical Analysis section.

### 5.7 ECG Processing

Heart rate was derived from ECG recordings acquired using the OpenBCI recording system. R-peaks were detected using the two-average detector implemented in the Python HeartPy/py-ecg-detectors package, and RR intervals were calculated as the time difference between successive R-peaks. Instantaneous heart rate was computed from the RR intervals and expressed in beats per minute (bpm). To obtain a continuous estimate of heart rate over time, mean heart rate was calculated within a 30-s sliding window advanced in 0.1-s increments. Heart-rate values were z-scored relative to each participant’s baseline period by subtracting the mean heart rate across the entire baseline phase and dividing by the corresponding baseline standard deviation. The resulting trajectories were aligned across experimental phases for group-level visualization. For participant-level statistical analyses, z-scored heart-rate values were averaged across all valid windows within each experimental phase to obtain one mean heart-rate value per participant and phase. Phase-related differences were evaluated according to the primary and secondary comparison framework described in the Statistical Analysis section.

### 5.8 EMG Processing

Swallow EMG was acquired during all three experimental phases; however, swallowing-related activity was analyzed only during the baseline and cephalic phases. Swallow-related EMG activity was quantified from the submental EMG channel. Raw EMG signals, sampled at 500 Hz, were bandpass filtered between 20 and 230 Hz using a fourth-order zero-phase Butterworth filter to isolate muscle activity and reduce low-frequency drift. Line-noise contamination was attenuated using a 60-Hz notch filter. The filtered EMG signal was then full-wave rectified and low-pass filtered at 2 Hz using a fourth-order zero-phase Butterworth filter to generate an EMG envelope representing swallowing-related muscle activation. Individual swallow events were identified during the baseline and cephalic phases using a peak-detection algorithm (scipy.signal.find peaks), with peaks required to exceed a predefined amplitude threshold and to be separated by a minimum interval of 10 s to reduce multiple detections of the same swallow. Detected swallow times were used to derive a time-resolved swallow-rate measure using a 3-min sliding window advanced in 1-min steps. Swallow-rate values were z-scored relative to each participant’s baseline period by subtracting the mean swallow rate across the entire baseline phase and dividing by the corresponding baseline standard deviation. The resulting trajectories were aligned across the baseline and cephalic phases for group-level visualization. For participant-level statistical analyses, z-scored swallow-rate values were averaged across all valid windows within each phase to obtain one mean swallow-rate value per participant and phase. Phase-related differences were evaluated according to the primary comparison framework described in the Statistical Analysis section.

### 5.9 Multiorgan Analysis

#### 5.9.1 Gastric–Cortical Phase–Amplitude Coupling

Time-varying gastric–cortical phase–amplitude coupling (PAC) was quantified using a Bayesian state-space framework following the approach described by Perley and Coleman [46]. This framework models coupling as a continuously evolving latent process, enabling time-resolved estimation of gastric–cortical interactions throughout the experimental period.

Gastric electrical activity (EGG) was bandpass filtered within the gastric slow-wave frequency range (0.03–0.07 Hz) using a fourth-order zero-phase Butterworth filter, and instantaneous gastric phase was extracted using the Hilbert transform. EEG signals were separately bandpass filtered within the theta (4–7 Hz), alpha (8–13 Hz), and beta (13–30 Hz) frequency bands using the same filtering approach, and instantaneous amplitude envelopes were obtained using the Hilbert transform. Within each cortical region, amplitude envelopes were summarized as the median across constituent electrodes. Gastric phase and regional cortical amplitude were subsequently down-sampled to a common 1-Hz time base and matched in duration before estimation of time-varying coupling.

The Bayesian model describes instantaneous EEG amplitude as a function of gastric phase using time-varying sinusoidal modulation coefficients that evolve according to a Gaussian random-walk state-space model. This formulation permits coupling strength to vary smoothly over time without imposing discrete transitions between fixed analysis windows. Model parameters and latent coefficient trajectories were estimated using a state-space Gamma generalized linear model implemented in JAX. Posterior uncertainty was approximated using a Laplace approximation, with a soft expectation–maximization (EM) procedure used to estimate the latent state trajectory and covariance structure. Samples were drawn from the resulting Gaussian approximation to the posterior distribution of the coefficient trajectories.

Sampled coefficient trajectories were transformed into time-resolved mutual information (MI) trajectories representing the estimated strength of gastric phase–cortical amplitude coupling. The posterior median MI trajectory was used as the primary time-resolved estimate, with the 2.5th and 97.5th percentiles of posterior MI samples providing 95% credible intervals. Maximum a posteriori (MAP) MI trajectories were additionally generated as complementary estimates of time-varying coupling.

Two complementary subject-level metrics were derived from the posterior median MI trajectories. First, mean MI was calculated for each experimental phase to quantify average gastric–cortical coupling strength. Second, positive excess MI area under the curve (AUC) was calculated by subtracting each participant’s region-specific mean baseline MI from the corresponding MI trajectory, retaining only positive deviations, and integrating these deviations over time using the trapezoidal rule. Positive excess AUC therefore quantified the cumulative magnitude and duration of above-baseline coupling elevations. The same participant- and region-specific baseline reference was applied across baseline, cephalic, and ingestion phases.

Whole-head PAC metrics were calculated by averaging the five regional estimates (frontal, parietal, left temporal, right temporal, and occipital) within each participant and experimental phase, with equal weighting across regions. Whole-head analyses required valid estimates from all five cortical regions. Mean MI and positive excess AUC were aggregated separately across regions rather than being recomputed from a pooled cortical amplitude envelope. For regional analyses, cephalic-phase changes in mean MI and positive excess AUC were calculated by subtracting each participant’s baseline value from the corresponding cephalic-phase value before group-level statistical comparisons and visualization.

Dynamic PAC analyses were performed independently for each participant, cortical region, frequency band, and experimental phase (baseline, cephalic, and ingestion). For time-resolved whole-head visualization, regional posterior median MI trajectories were first baseline-centered using each participant’s region-specific mean baseline MI, interpolated onto a common 5-s time grid within each 15-min phase, and averaged across the five regions within each participant. Group-level trajectories were then summarized as the mean *±* standard error of the mean (SEM) across participants, without extrapolation beyond the available temporal support.

Primary statistical comparisons evaluated baseline–cephalic changes in mean MI and positive excess AUC using paired *t*-tests. Whole-head comparisons were performed separately for theta, alpha, and beta frequency bands. Regional comparisons were conducted independently for each coupling metric and frequency band, with *p*-values corrected across the five cortical regions using the Benjamini–Hochberg false discovery rate (FDR) procedure. Comparisons involving ingestion were treated as secondary analyses, with Holm adjustment applied within each coupling measure.

To assess whether significant regional theta positive excess AUC findings were driven by individual participants, leave-one-participant-out sensitivity analyses were performed. Each participant was excluded in turn, and baseline–cephalic comparisons were repeated with FDR correction across the five cortical regions. The persistence of significant effects across these analyses was used to assess robustness to individual observations.

#### 5.9.2 Physiological Correlates of Subjective Appetite

Associations between cephalic-phase physiological responses and subjective appetite were examined using baseline-to-cephalic changes in physiological activity. For each physiological measure, change was calculated as the mean value during the cephalic phase minus the corresponding baseline mean. Changes in self-reported hunger and satiety from before baseline to immediately following the cephalic phase were assessed using paired-samples *t*-tests. For analyses examining associations with physiological responses, changes in hunger and satiety and post-cephalic craving ratings were treated as ordinal outcomes and analyzed using ordinal logistic regression.

Gastric associations were assessed using changes in gastric myoelectric power. Cortical associations were assessed using changes in normalized EEG spectral power within the theta (4–7 Hz), alpha (8–13 Hz), and beta (13–30 Hz) frequency bands. Band-specific power was normalized to total spectral power from 1–30 Hz. Cortical measures were calculated separately across frontal, parietal, left temporal, right temporal, and occipital regions by averaging normalized spectral power across electrodes within each region. A log-transformed alpha/beta power ratio was additionally calculated to characterize the relative balance between alpha and beta activity and capture coordinated changes across these opposing spectral responses. The log(alpha/beta) ratio was calculated separately during the baseline and cephalic phases, followed by calculation of the cephalic-minus-baseline change score.

For regional cortical–subjective analyses, extreme predictor values were identified separately for each cortical measure and region using a prespecified 3*×*IQR criterion. Values falling below *Q*_1_ *−* 3 *×* IQR or above *Q*_3_ + 3 *×* IQR were excluded prior to ordinal logistic regression. Regional cortical *p*-values were corrected across the five cortical regions using the Benjamini–Hochberg false discovery rate (FDR) procedure, according to the multiple-comparison framework described in the Statistical Analysis section.

To determine whether gastric and cortical responses independently contributed to subjective craving, separate multivariable ordinal logistic regression models were fitted for each of the five cortical regions. Cephalic-minus-baseline changes in gastric myoelectric power and the corresponding regional log(alpha/beta) ratio were entered simultaneously as predictors of craving. Both continuous predictors were standardized within each regional model such that coefficients and odds ratios reflected a one-standard-deviation increase in the corresponding physiological predictor. Multivariable analyses included all participants with complete gastric, cortical, and craving data (*n* = 40), with no additional outlier-based exclusions. The *p*-values associated with the cortical log(alpha/beta) predictor were corrected across the five regional models using the Benjamini–Hochberg FDR procedure.

### 5.10 Statistical Analysis

Statistical analyses were structured around the primary objective of characterizing physiological changes associated with the cephalic phase of digestion. Accordingly, the baseline-to-cephalic comparison was specified as the primary planned comparison. Comparisons involving the ingestion phase (cephalic-to-ingestion and baseline-to-ingestion) were considered secondary and were used to assess the persistence or evolution of physiological responses following food consumption.

For physiological analyses comparing all three experimental phases, overall phase effects were assessed using repeated-measures analysis of variance (ANOVA) in participants with complete data across baseline, cephalic, and ingestion phases. The primary baseline-to-cephalic comparison was evaluated using a paired-samples *t*-test in all participants with available data for these two phases. The two secondary pairwise comparisons (cephalic-to-ingestion and baseline-to-ingestion) were corrected jointly for multiple comparisons using the Holm procedure. Effect sizes for paired comparisons were quantified using Cohen’s *d_z_*. Statistical tests were two-sided, and *p <* 0.05 was considered statistically significant.

For regional analyses, multiple-comparison correction was performed separately within each frequency band and outcome measure. Primary baseline-to-cephalic comparisons were corrected across the five cortical regions using the Benjamini–Hochberg FDR procedure. The two secondary phase comparisons (cephalic- to-ingestion and baseline-to-ingestion) were treated as a single family and corrected across the 10 resulting tests (five cortical regions *×* two phase comparisons) using the Benjamini–Hochberg procedure.

For visualization of time-resolved physiological measures, participant-level trajectories were aligned by experimental phase and averaged across participants at each time point. Group-level trajectories are presented as the mean *±* standard error of the mean (SEM), unless otherwise specified.

## Acknowledgements

The authors acknowledge the support of the Koret Human Neurosciences Community Laboratory and thank Sophia Shen and Suchita Rao for their assistance with experimental data collection.

## Funding

This work was supported by the Stanford Interdisciplinary Graduate Fellowship (to Y.M.K.), Stanford Bio-X (to T.P.C.), the Wu Tsai Neurosciences Institute (to T.P.C.), the Yu Faculty Scholar funds from the Stanford School of Engineering (to T.P.C.), and the National Institutes of Health (NIH) grant 1U01DK140939 (to T.P.C.).

## Competing interests

The authors declare no competing interests.

## Notes

### Competing Interest Statement

The authors have declared no competing interest.

### Summary of Updates

The figures were duplicated, thus we removed the figures that appear at the end of the preprint.

